# Germline TFAM levels regulate mitochondrial DNA copy number and mutant heteroplasmy in *C. elegans*

**DOI:** 10.1101/2023.01.28.526030

**Authors:** Aaron Z.A. Schwartz, Jeremy Nance

## Abstract

The mitochondrial genome (mtDNA) is packaged into discrete protein-DNA complexes called nucleoids. mtDNA packaging factor TFAM (mitochondrial transcription factor-A) promotes nucleoid compaction and is required for mtDNA replication. Here, we investigate how changing TFAM levels affects mtDNA in the *Caenorhabditis elegans* germ line. We show that increasing germline TFAM activity boosts mtDNA number and significantly increases the relative proportion of a selfish mtDNA mutant, *uaDf5*. We conclude that TFAM levels must be tightly controlled to ensure appropriate mtDNA composition in the germ line.

## Description

The mitochondrial transcription factor-A (TFAM) plays essential roles in regulating mtDNA copy number, compacting nucleoids, and replicating/transcribing mtDNA (Garrido et al. 2003; Lewis et al. 2016; Fu et al. 2020). Global reduction of TFAM activity has a conserved effect on mtDNA copy number in metazoans: genetic knockdown severely reduces mtDNA levels in mammals, fish, flies, and cell culture systems (Larsson et al. 1998; Kanki et al. 2004; Matsushima et al. 2004; Otten et al. 2020; Wang et al. 2021). The effects of TFAM overexpression on mtDNA levels are less clear. Various studies in cell culture demonstrate that TFAM overexpression can be both sufficient (Kanki et al. 2004; Matsushima et al. 2004), and insufficient (Maniura-Weber et al. 2004) to drive increases in mtDNA. The situation *in vivo* is similar, as two independent studies in *Drosophila* saw no net effect of TFAM overexpression on mtDNA levels (Matsuda et al. 2013; Cagin et al. 2015), whereas TFAM overexpression in mice was sufficient to drive mtDNA expansion above normal levels (Ekstrand et al. 2004). Previously, we and others showed that global reduction in TFAM activity has a profound negative impact on mtDNA levels in *C. elegans*, as expected (Sumitani et al. 2011; Lin et al. 2016; Schwartz et al. 2022). Here we ask the converse: is germline overexpression of the worm *TFAM* homolog, *hmg-5*, sufficient to increase mtDNA levels *in vivo*?

To overexpress TFAM (encoded by the *hmg-5* gene) in the germ line, we used regulatory elements from the endogenous *glh-1* gene, which encodes a highly expressed germline-specific protein (Marnik et al. 2019; Goudeau et al. 2021). We employed two strategies to express untagged TFAM at the *glh-1* locus: the viral 2A self-cleaving peptide system, which results in the production of two peptides via ribosomal skipping during translation (**Fig. 1A, left)**; and the *C. elegans SL2* trans-splicing element (derived from the operonic gene *rla-1*), which causes the nascent transcript to be spliced into two independently translated mRNAs (**Fig. 1A, right)** (Nance and Frokjaer-Jensen 2019). We used CRIPSR/Cas9 genome engineering to insert either *T2A::TFAM* or *SL2::TFAM* at the 3’ end of the endogenous *glh-1* protein coding sequence (**Fig. 1A**). To avoid known loss of functionality due to the presence of C-terminal tags on TFAM (Schwartz et al. 2022), we expressed TFAM untagged.

**Figure 1.**
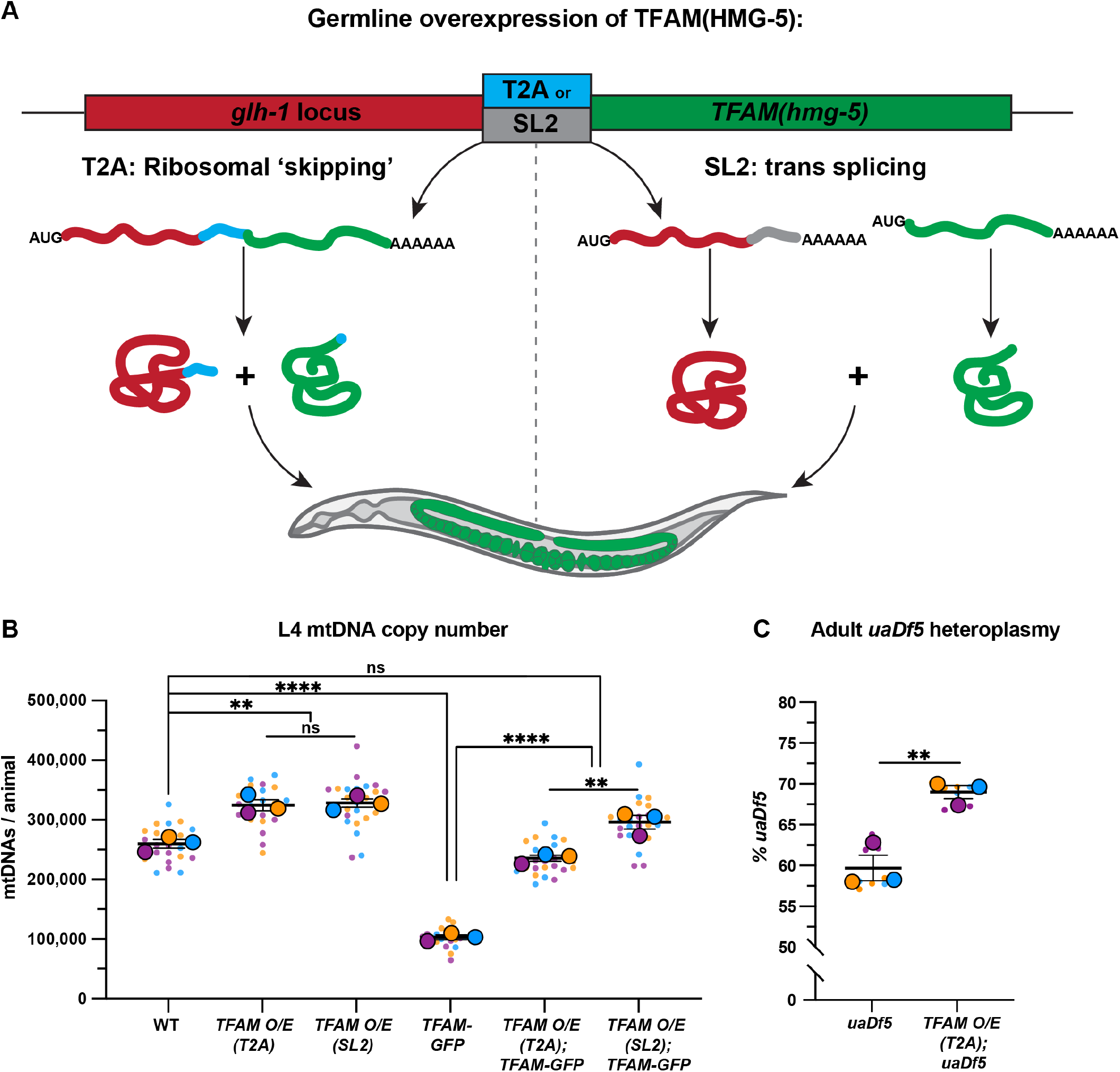
Germline TFAM overexpression modulates mtDNA copy number and heteroplasmy. (**A**) Schematic of strategies to increase germline TFAM levels by overexpression at the endogenous *glh-1* locus. (**B**) Quantification of total mtDNA copy number from whole L4 larvae by qPCR in wild-type, *TFAM-O/E (T2A), TFAM-O/E (SL2), TFAM-GFP, TFAM-O/E (T2A); TFAM-GFP*, and *TFAM-O/E (SL2); TFAM-GFP* animals. (**C**) Quantification of *uaDf5* heteroplasmy in *uaDf5* and *TFAM-O/E (T2A); uaDf5* adults. Small dots are data points from individual L4 worms in (B), and technical ddPCR replicates in (C), from each of three color-coded biological replicates; the mean from each replicate is shown as a larger circle, the mean of means as a horizonal line, and the S.E.M as error bars. n.s., not significant (*p*> 0.05), \*\**p* ≤ 0.01, **** *p* ≤ 0.0001, unpaired two-tailed Student’s *t*-test.

To determine the effect of overexpressing TFAM on mtDNA levels, we measured total mtDNA in whole L4 larvae by quantitative PCR (qPCR), as ∼90% of total *C. elegans* mtDNA is derived from the germline (Tsang and Lemire 2002; Bratic et al. 2009). Both means of overexpressing TFAM resulted in a significant increase in mtDNA levels (∼25-30%) (**Fig. 1B**). Conversely, as reported previously, we found that animals homozygous for a hypomorphic *TFAM/hmg-5* allele (endogenously tagged *TFAM-gfp)* had severe mtDNA copy number defects in L4 larvae (**Fig. 1B**) (Schwartz et al. 2022). This defect in mtDNA number could be rescued by overexpressing TFAM in the *TFAM-gfp* hypomorphic background (**Fig. 1B**). Together, these results demonstrate that excess TFAM is sufficient to drive an increase in mtDNA levels in the *C. elegans* germ line.

We next sought to determine if overexpressing TFAM affected mtDNA quality. To accomplish this, we used a strain containing the *uaDf5* mtDNA deletion, which removes 3.1kb of the 13.8kb mitochondrial genome (Tsang and Lemire 2002). *uaDf5* deletes essential genes and therefore must exist in heteroplasmy with complementing wild-type mtDNA. Because *uaDf5* mtDNA has a selfish replicative advantage over wild-type mtDNAs (Gitschlag et al. 2020; Yang et al. 2022; Schwartz et al. 2022), we hypothesized that TFAM overexpression would increase the proportion of *uaDf5* mtDNA relative to wild-type mtDNA. Indeed, *TFAM-O/E (T2A); uaDf5* adult animals contained a significantly higher proportion of mutant mtDNAs (∼70%) compared to *uaDf5* controls (∼60%) (**Fig. 1C**). This finding suggests that increasing mtDNA levels by overexpressing TFAM favors an even further expansion of *uaDf5* mutant genomes over wild type mtDNAs compared to controls expressing normal TFAM levels.

## Methods

### Worm culture and strains

*C. elegans* strains were maintained at 20°C on nematode growth medium plates seeded with *Escherichia coli* OP50 as previously described (Brenner 1974). A list of all strains used/generated in the study is available in the *<u>Strain Table</u>* below.

### Mitochondrial DNA quantification

Quantification of total mtDNA copy number of whole L4 larvae via qPCR was performed exactly as previously described (Schwartz et al. 2022). For *uaDf5* heteroplasmy measurement, 60-100 whole adult animals were pooled in 60-100µL of worm lysis buffer [50 mM KCl, 10 mM Tris-HCl (pH 8.0), 2.5 mM MgCl_2_, 0.45% IGEPAL (Sigma I8896)] in a screw cap 1.5mL microfuge tube, flash frozen at -80°C for 15 minutes, and lysed in a heating block at 60°C for 1 hour followed by 15 minutes at 95°C. Adult worm lysates were diluted 1000X and droplet digital PCR (ddPCR) quantification of *uaDf5* and WT mtDNA was performed exactly as previously described (Schwartz et al. 2022).

### Plasmid construction

Plasmid pJN651 (*glh-1::SL2::YFP::PH::glh-1 3’UTR)* was constructed by replacing *GFP* in pDU92 (*glh-1::SL2::GFP::glh-1 3’UTR)* by Gibson assembly (Gibson, Young et al. 2009).

### CRISPR/Cas9 genome editing

CRISPR/Cas9-mediated genome editing was performed as described previously (Paix et al. 2017; Schwartz et al. 2022). Construction of *glh-1*(*xn127[glh-1::SL2::hmg-5])* required two steps. First, *glh-1(xn81[glh-1::SL2::yfp-PH]*) was generated using pJN651 as a PCR template to amplify *SL2::yfp-PH* with ∼35 bps of homology for insertion at the C-terminus of endogenous *glh-1*. Second, for the generation of *glh-1*(*xn127[glh-1::SL2::hmg-5])*, N2 genomic DNA was used as a PCR template to amplify *hmg-5* with ∼35bp of homology to replace *yfp-PH* by CRISPR at the *glh-1(xn81[glh-1::SL2::yfp-PH]*) locus. To generate *glh-1*(*xn167[glh-1::T2A::hmg-5])*, a single stranded oligonucleotide template was used to swap *SL2 for T2A* via CRISPR at the *glh-1*(*xn127[glh-1::SL2::hmg-5])* locus. Sequence files for all insertions are available upon request. All gRNA sequences are found in the *<u>Sequences Table</u>* below

### Statistical analysis and reproducibility

All statistical analysis was performed using GraphPad Prism 9 software. For all data, unpaired two-tailed Student’s *t*-tests were performed, and where applicable no corrections for multiple comparisons were made to avoid type II errors (Armstrong 2014). Data in graphs are shown as Superplots (Lord et al. 2020). Three biologically independent experiments were performed for all experiments and the arithmetic means of biological replicates were used for statistical analysis.

## Reagents

***Strain Table:***

| Strain | Genotype | Source |
| --- | --- | --- |
| FT2168 | *naSi2 [mex-5p::mCherry-H2B::nos-2 3'UTR, unc-119(+)] II* | Nance Lab (*naSi2* transgene from (Roy et al. 2018) outcrossed twice |
| FT2278 | *glh-1(xn167[glh-1::T2A::hmg-5]) I; naSi2 [mex-5p::mCherry-H2B::nos-2 3'UTR, unc-119(+)] II* | Nance Lab (This study) |
| FT2217 | *glh-1(xn127[glh-1::SL2::hmg-5]) I; naSi2 [mex-5p::mCherry-H2B::nos-2 3'UTR, unc-119(+)] II* | Nance Lab (This study) |
| FT2133 | *xnSi45 [mex-5p::mCherry-MOMA-1::nos-2 3'UTR; unc-119(+)] II; hmg-5(xn107[hmg-5-gfp]) IV* | Nance Lab (This study) |
| FT2326 | *glh-1(xn167[glh-1::T2A::hmg-5]) I; xnSi45 [mex-5p::mCherry-MOMA-1::nos-2 3'UTR; unc-119(+)] II; hmg-5(xn107[hmg-5-gfp]) IV* | Nance Lab (This study) |
| FT2285 | *glh-1(xn127[glh-1::SL2::hmg-5]) I; xnSi45 [mex-5p::mCherry-MOMA-1::nos-2 3'UTR; unc-119(+)] II; hmg-5(xn107[hmg-5-gfp]) IV* | Nance Lab (This study) |
| FT2283 | *glh-1(sam24[glh-1::gfp::3Xflag]) I; xnIs510 [ehn-3::mCherry-PH, unc-119(+)] II; uaDf5 / + mtDNA* | Nance Lab (This study) |
| FT2409 | *glh-1(xn167[glh-1::T2A::hmg-5]) I; xnIs510 [ehn-3::mCherry-PH, unc-119(+)] II; uaDf5 / +* | Nance Lab (This study) |
| FT1917 | *glh-1(xn81[glh-1::SL2::YFP-PH]) I; naSi2 [mex-5::mCherry-H2B::nos-2 3' UTR, unc-119(+)] II* | Nance Lab (This study) |

***Sequences Table***

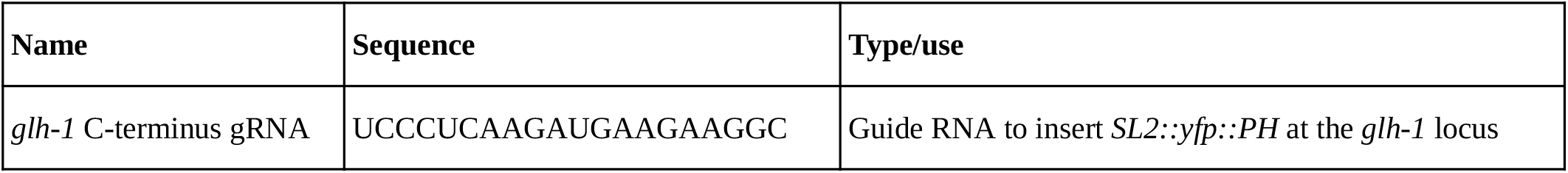
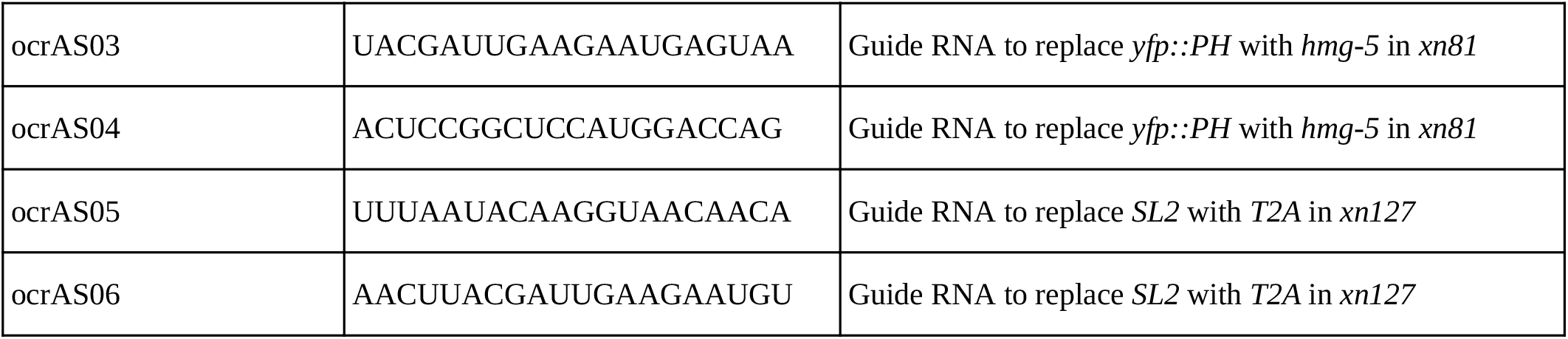

## Author Contributions

Aaron Z.A. Schwartz: writing - original draft, conceptualization, formal analysis, investigation. Jeremy Nance: writing - original draft, supervision, resources.

